# HPF1-dependent histone ADP-ribosylation triggers chromatin relaxation to promote the recruitment of repair factors at sites of DNA damage

**DOI:** 10.1101/2021.08.27.457930

**Authors:** Rebecca Smith, Siham Zentout, Catherine Chapuis, Gyula Timinszky, Sébastien Huet

## Abstract

PARP1 activity is regulated by its cofactor HPF1. The binding of HPF1 on PARP1 controls the grafting of ADP-ribose moieties on serine residues of proteins nearby the DNA lesions, mainly PARP1 and histones. However, the impact of HPF1 on DNA repair regulated by PARP1 remains unclear. Here, we show that HPF1 controls both the number and the length of the ADP-ribose chains generated by PARP1 at DNA lesions. We demonstrate that HPF1-dependent histone ADP-ribosylation, rather than auto-modification of PARP1, triggers the rapid unfolding of the chromatin structure at the DNA damage sites and promotes the recruitment of the repair factors CHD4 and CHD7. Together with the observation that HPF1 contributes to efficient repair both by homologous recombination and non-homologous end joining, our findings highlight the key roles played by this PARP1 cofactor at early stages of the DNA damage response.

## INTRODUCTION

Poly(ADP-ribose) Polymerase 1 (PARP1), which belongs to the diphtheria toxin-like family of ADP-riboyltransferases, is the founding member of a large family of enzymes that regulate a number of different cellular processes. PARP1 itself plays pivotal functions in DNA repair, chromatin folding and gene transcription^1,2^. As part of its role in the DNA damage response (DDR), PARP1 detects both single and double strand breaks^3^ through its N-terminal DNA-binding domain consisting of three zinc fingers modules^4^. The binding of this domain to DNA breaks triggers the catalytic activity of the C-terminal domain via a complex allosteric mechanism^5^. Once activated, PARP1 utilizes NAD+ to polymerize ADP-ribose (ADPr) chains on target proteins, with the major targets being histones and PARP1 itself ^6,7^.

Early research into PARP1 focused on its role as a discrete enzyme, capable of catalyzing the addition of ADP-ribose chains alone. However, recent studies identified a key co-factor, Histone Poly(ADP-ribosylation) Factor 1 (HPF1), which is required for targeting ADPr chains on specific residues^8^. Indeed, HPF1 binding to the C-terminus of PARP1 creates a joint catalytic site that is essential to ADP-ribosylate serines^9^,which are the main residues to be modified by ADPr in the context of the DDR^10^. The loss of HPF1 therefore has several effects; firstly, auto-modification of PARP1 shifts from occurring on serines to primarily to glutamic and aspartic acids and secondly, trans ADP-ribosylation of histones is suppressed^11^. Moreover, *in vitro*, HPF1 is not only required for targeting ADPr to specific residues, but it also controls the rate of polymerization, favoring mono-ADPr modifications over poly-ADPr chains^12^

The major findings reported over the last years have allowed to considerably improve our understanding of the molecular mechanisms underlying the control of ADP-ribosylation signaling by the PARP1/HPF1 axis. Nevertheless, the exact impact of HPF1 on cellular functions known to be regulated by PARP1 remains unclear. A role for HFP1 in DNA repair has been hinted at as HPF1-deficiency led to cell hypersensitivity to DNA-damaging agents^8^. Therefore, in this report, we aimed to further investigate how HPF1 could regulate ADP-ribosylation-dependent steps of the DDR. We show that HPF1 is recruited to DNA lesions via its binding to the C-terminal residues of PARP1 and that it controls both the number and length of ADPr chains at sites of damage. We also establish that HPF1-dependent histone ADP-ribosylation, rather than PARP1 auto-modification, is a major trigger of the early chromatin unfolding that occurs in the vicinity of the DNA lesions, thus facilitating the recruitment of subsequent repair factors. Therefore, we demonstrate that HPF1 is a central player at early stages of the DDR and that its role in the regulation of chromatin structure contributes to efficient DNA repair.

## RESULTS

### HPF1 recruitment to sites of DNA damage relies on interaction with the C-terminus of PARP1

HPF1 interacts with the C-terminal end of PARP1, the last two residues L1013/W1014 of the latter being essential for this interaction^9,13^. Nevertheless, the contribution of this interaction to the rapid recruitment of HPF1 at DNA lesions^8^ remains unclear since recent findings suggest that HPF1 and PARP accumulate to sites of damage independently from each other^14^. To investigate this question further, we first quantified the recruitment of both HPF1 and PARP1 to sites of damage induced by laser-microirradiation in U2OS *PARP1*^*KO*^ cells co-expressing mCherry-PARP1 and GFP-HPF1 (Figure 1A,B). While the recruitment of both proteins peaked within 10 sec, we observed that PARP1 accumulation to sites of damage was much stronger than HPF1. This difference in the relative amounts of the two proteins accumulating at DNA lesions is in line with the *in vitro* data suggesting that HPF1 can exert its regulatory role on PARP1 even at low relative molarity^15^. Interestingly, HPF1 dissipated from the damage slower than PARP1 (Fig 1C), indicating that the HPF1/PARP1 molarity ratio increases progressively at DNA lesions during the first 10 minutes following damage induction.

**Figure 1:**
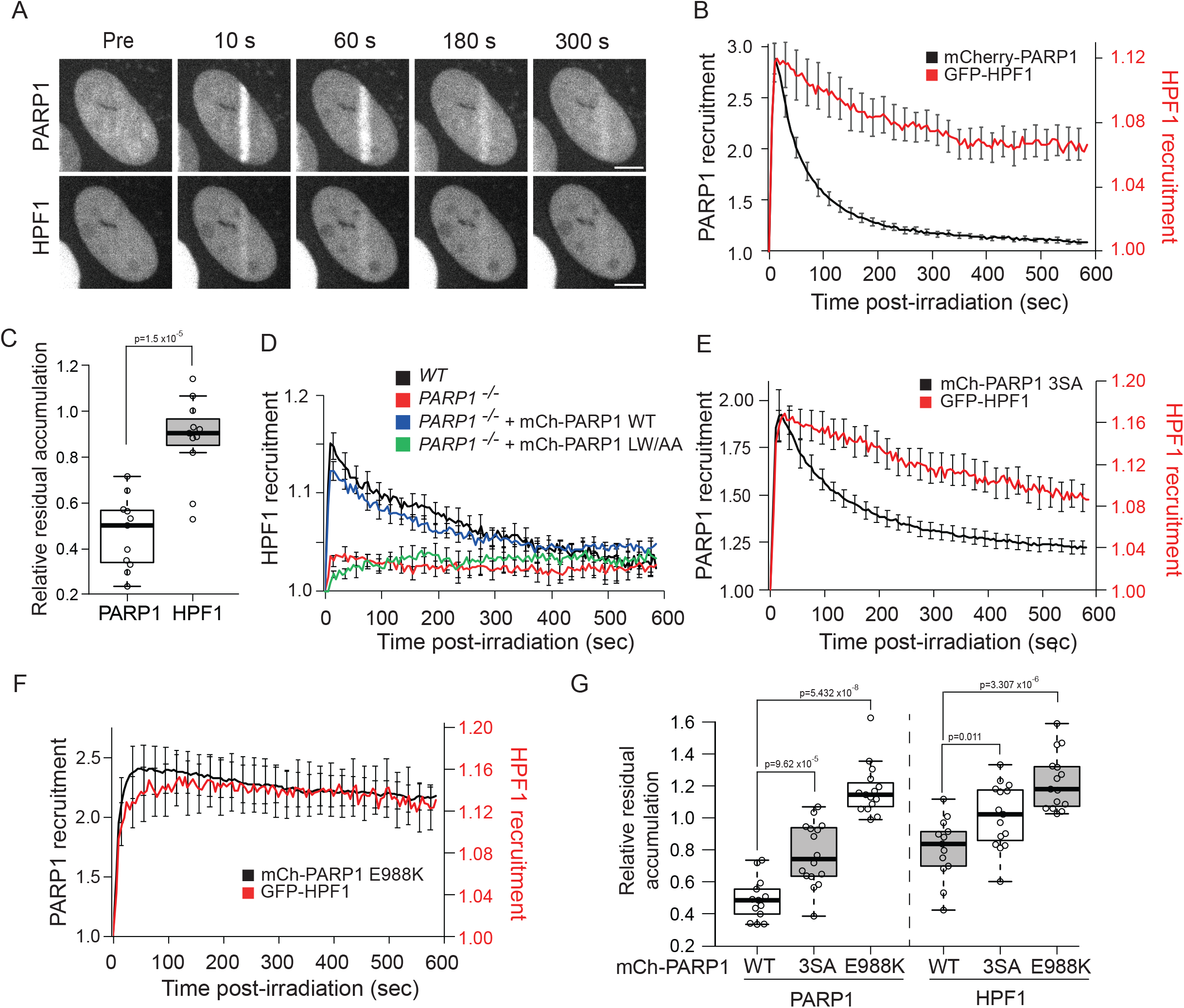
HPF1 recruitment to sites of damage relies on interaction with the C-terminus of PARP1. (**A**) Representative images of mCherry-PARP1 and GFP-HPF1 recruitment to sites of DNA damage induced by laser irradiation, in *PARP1*^*KO*^ cells. Scale bar, 5 μm. (**B**) Recruitment kinetics of mCherry-PARP1 (black) and GFP-HPF1 (red) to sites of DNA damage. (**C**) To assess the relative release kinetics of mCherry-PARP1 and GFP-HPF1, the time *t*_*1/2*_ at which half of PARP1-WT has been released compared to peak accumulation was first estimated from the mean curve shown in B. Then, the amount of the different proteins is measured at *t*_*1/2*_ for each individual cell and normalized to peak accumulation to estimate the relative residual accumulation. (**D**) Recruitment kinetics of GFP-HPF1 to sites of DNA damage in *WT* or *PARP1*^*KO*^ cells expressing mCherry-taggedPARP1 WT or PARP1 L1013A/W1014A (PARP1-LW/AA). (**E, F**) Recruitment kinetics of (**E**) mCherry-PARP1-3SA (black) or (**F**) mCherry-PARP1-E988K (black) and GFP-HPF1 (red) to sites of DNA damage. (**G**) With the same approach as for panel C, the relative residual accumulation of mCherry tagged PARP1 mutants and GFP-HPF1 was estimated from the curves shown in E and F.

In agreement with previous results^8^, we found that PARP1 deficiency nearly fully suppressed HPF1 recruitment to DNA lesions, the presence of PARP2 being unable to compensate for PARP1 loss despite the known interaction between HPF1 and PARP2 (Fig 1D and Fig S1A). HPF1 recruitment in *PARP1*^*KO*^ cells was rescued upon re-expression of a wild-type version of PARP1 (PARP1-WT) but not in the presence of PARP1 mutated at residues L1013A/W1014A (PARP1-LW/AA) (Fig 1D). These data indicate that the interaction of HPF1 with these last two PARP1 residues is critical for HPF1 accumulation to sites of damage. We also observed that PARP1 tagged at its C-terminal end is unable to rescue HPF1 recruitment when expressed in *PARP1*^*KO*^ cells, therefore, this tagging strategy should be avoided when assessing PARP1 behavior at sites of damage (Fig S1B).

Beside its recruitment to sites of damage, we also wondered whether HPF1 release could be regulated by the mobilization of PARP1 from this area. Auto-ADP-ribosylation of PARP1 is a key regulator of its release from the DNA lesions^16^. Therefore, we analyzed HPF1 release in *PARP1*^*KO*^ cells re-expressing PARP1 mutants displaying impaired auto-ADP-ribosylation (Fig S1B,C) due to either suppressed catalytic activity (PARP1 E988K) or mutations of the main serine residues targeted for ADP-ribosylation on PARP1 (PARP1-S499/507/519A, PARP1-3SA). As previously shown^16–18^, both mutants were retained longer at sites of damage compared to PARP1-WT and we observed that HPF1 release kinetics mirrored the relative dissipation speeds of the different PARP1 mutants (Fig 1E-G). These findings highlight that the transient accumulation of HPF1 at damage sites is tightly controlled by PARP1 during both the accumulation and the release phases.

### HPF1 controls the number and length of ADP-ribose chains at sites of damage

HPF1 controls both the targeting of ADP-ribose (ADPr) chains on specific residues and the rate of ADP-ribosylation^11,15^. Therefore, we wondered what the overall impact of the loss of HPF1 on ADP-ribosylation signaling was at sites of damage by analyzing the recruitment kinetics of two different ADPr-binding domains. Firstly, the macrodomain of macroH2A1.1, which associates with mono-ADPr or the terminal residue of poly-ADPr chains (^19,20^, Fig 2A), was used as a proxy for the number of ADPr chains at sites of damage. Secondly, the WWE domain of RNF146, which binds at the interface between two monomers along poly-ADPr chains, was used to estimate the total amount of ADPr (^21^, Fig 2A). Both ADPr-binding domains showed similar behavior upon laser irradiation: an initial peak of recruitment within the first 30 s post damage followed by a stable or slowly decreasing plateau within the next 10 minutes (Fig 2B-E). These recruitment profiles suggest that ADP-ribosylation signaling starts with an early acute phase followed by a more sustained period lasting for several minutes after damage. Importantly, this sustained signaling period arises from a dynamic equilibrium between ADP-ribose polymerase and hydrolase activities since the acute inhibition of PARP1 during this period leads to the rapid removal of the ADPr chains from the sites of damage^22^.

**Figure 2:**
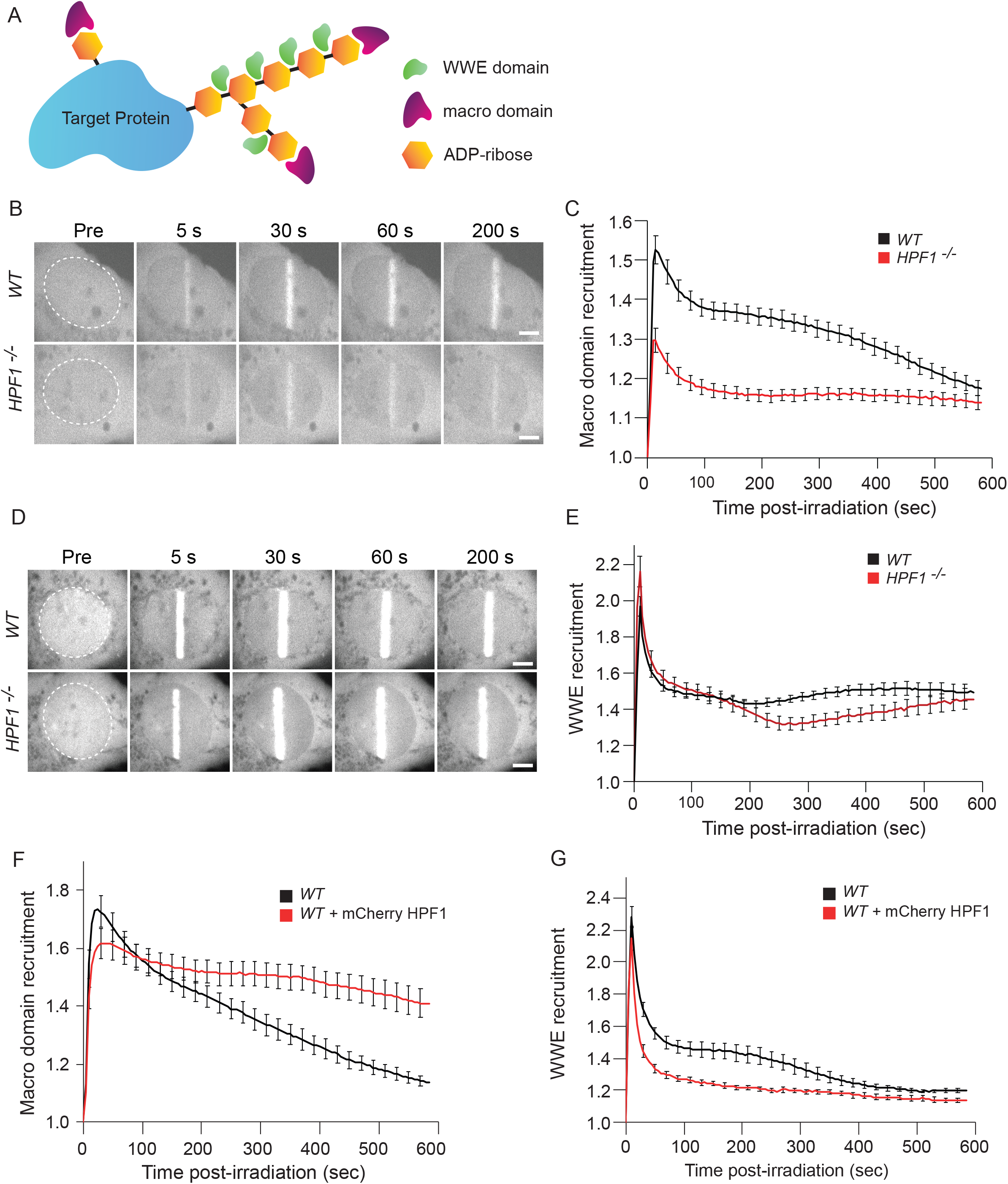
HPF1 regulates ADP-ribosylation signalling at sites of DNA damage. (**A**) Schematic representation of WWE and macrodomain recruitment on ADPr chains. (**B**) Representative images showing recruitment of the macrodomain of macroH2A1.1 to sites of DNA damage induced by laser irradiation in *WT* and *HPF1*^*KO*^ cells. Scale bar, 5 μm. (**C**) Recruitment kinetics of the macrodomain to sites of DNA damage in *WT* (black) and *HPF1*^*KO*^ (red) cells. (**D**) Representative images showing recruitment of the WWE domain of RNF146 to sites of laser induced DNA damage in *WT* and *HPF1*^*KO*^ cells. Scale bar, 5 μm. (**E**) Recruitment kinetics of the WWE domain to sites of DNA damage in *WT* (black) and *HPF1*^*KO*^ (red) cells. (**F, G)** Recruitment kinetics of (**F**) GFP-WWE or (**G**) GFP-macrodomain of mH2A1.1 recruitment kinetics at sites of DNA damage in U2OS *WT* cells overexpressing mCherry-HPF1 or not. Data are shown as mean ± SEM.

We found that HPF1 deficiency was associated with a strong reduction in macrodomain recruitment (Fig 2B,C). Similar defect was also observed in *PARP1*^*KO*^ cells expressing PARP1-LW/AA, which is unable to recruit HPF1 to DNA lesions, compared to those expressing PARP1-WT (Fig S2A,C). Therefore, the absence of HPF1 at sites of damage led to a decrease in the number of ADPr chains that are generated upon damage. Conversely, the loss of HPF1 had no major impact on WWE accumulation to sites of damage (Fig 2D,E), showing that the total amount of ADPr generated at sites of damage remained unchanged. This, together with the data regarding macrodomain recruitment, suggests that the fewer chains generated in the absence of HPF1 are longer. These findings agree with previous *in vitro* results showing that HPF1 is not only crucial to initiate the grafting of ADPr chains on a significant number of acceptor residues but also restrains ADPr chain length^12^.

Recent reports have also shown that modifying the relative molarities of HPF1 and PARP1 strongly impacts ADP-ribosylation activity *in vitro*^12,15^. Therefore, we wondered what the impact of the over-expression of HPF1 was, whose endogenous nuclear levels are 20 to 50 times lower than those of PARP1^8^, on ADP-ribosylation signaling at sites of damage. HPF1 overexpression had little influence on the initial peak of both macrodomain and WWE but perturbed the slower plateau phases (Fig 2F,G) with increased accumulation of macrodomain while WWE was reduced at sites of damage. Therefore, the excess of HPF1 appears to increase the amount of ADPr chains while reducing their lengths. These data show that the relative amounts of HPF1 versus PARP1 within the nucleus controls the characteristics of ADP-ribosylation signaling at sites of damage.

### HPF1 is a key regulator of early chromatin relaxation at sites of DNA damage

In addition to its role in signaling the presence of DNA lesions for repair effectors, ADP-ribosylation by PARP1 also triggers rapid chromatin relaxation in the vicinity of DNA breaks^23^, a process that facilitates the access to the lesions^22^. Therefore, we wondered whether HPF1 could also regulate this early chromatin remodeling process due to its regulatory role in ADP-ribosylation signaling. To assess this, we monitored the amount of chromatin relaxation in WT, *PARP1*^*KO*^, *HPF1*^*KO*^ and *PARP1*^*KO*^/*HPF1*^*KO*^ cells by a live-cell chromatin relaxation assay (Fig 3A,B). In this assay, a region of chromatin is highlighted by the local photoconversion of PAGFP fused to the histone H2B, which occurs simultaneously to damage induction by laser irradiation at 405 nm. The changes in the level of chromatin condensation at the sites of damage are estimated by measuring the thickness of the photoconverted line. Chromatin relaxation at sites of damage was nearly fully suppressed in the absence of PARP1. The loss of HPF1 also dramatically reduced chromatin relaxation, although not to the same degree as in *PARP1*^*KO*^ cells. The concomitant loss of PARP1 and HPF1 led to relaxation levels that were lower to those in *HPF1*^*KO*^ cells, showing that the residual relaxation seen in these cells remains PARP1-dependent. The defect in chromatin relaxation seen in *HPF1*^*KO*^ cells is the consequence of the absence of HPF1 at DNA lesions. Indeed, the expression in *PARP1*^*KO*^ cells of the PARP1-LW/AA mutant or C-terminally tagged PARP1, which both fail to recruit HPF1 to sites of damage, was unable to restore chromatin relaxation at the level measured in cells re-expressing PARP1-WT (Fig 3C, Fig S3A). Similarly, the expression of the HPF1 mutant D283A, that did not to recruit to damage (Fig S3B) due to impaired interaction with PARP1^9,13^, did not rescue chromatin relaxation in *HPF1*^*KO*^ cells (Fig 3D).

**Figure 3:**
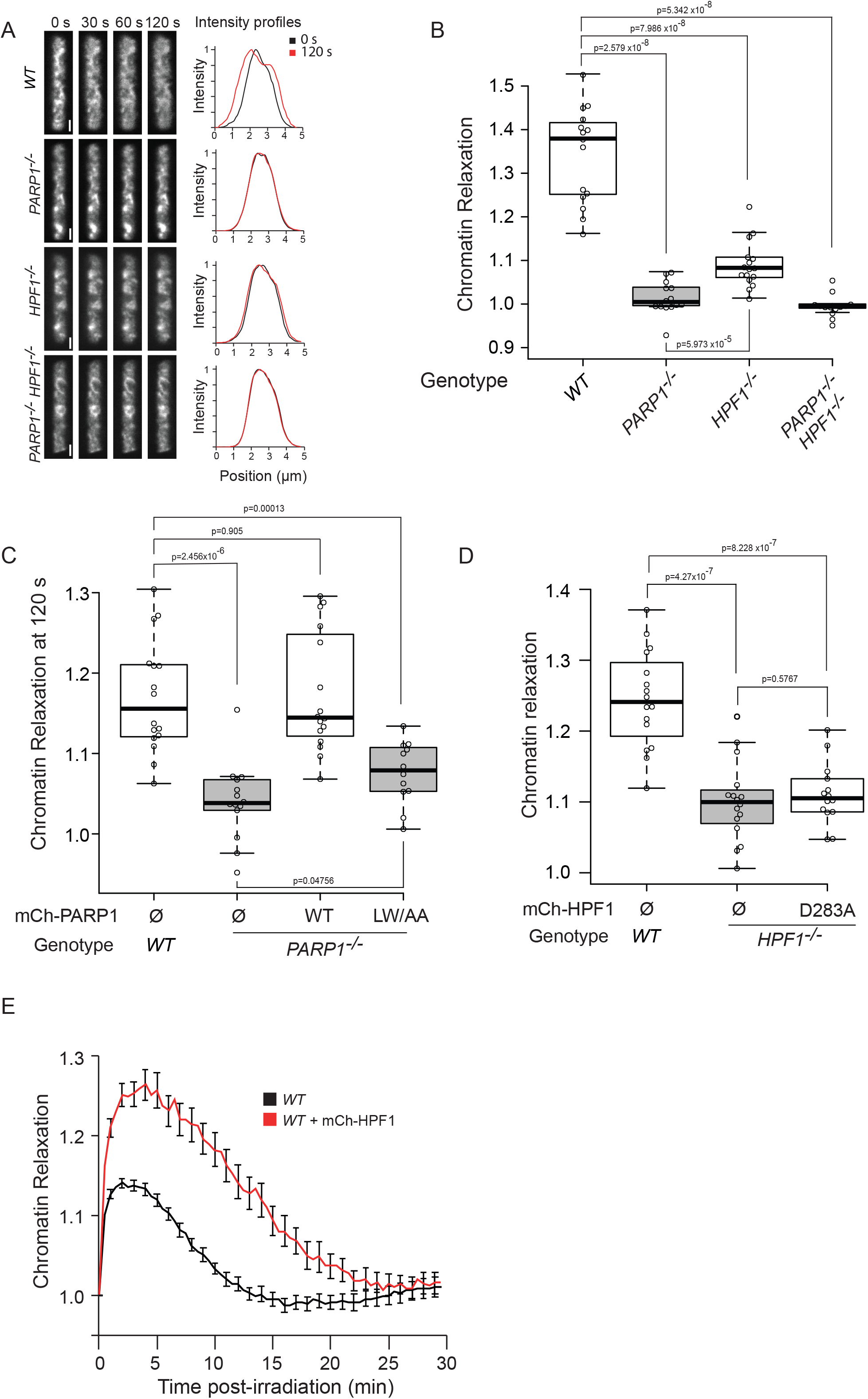
HPF1 promotes chromatin relaxation at sites of DNA damage. (**A**) Left: Confocal image sequences of the chromatin line area which got simultaneously damaged and photoconverted by irradiation at 405 nm in U2OS *WT*, *PARP1*^*KO*^, *HPF1*^*KO*^ and *PARP1/HPF1* double knockout cells expressing H2B-PAGFP. Scale bars, 2 μm. Right: Intensity profiles perpendicular to the irradiated lines (μm) at 0 s (black) and 120 s (red) after damage induction. The enlargement of the profile is due to the thickening of the photoconverted line consecutive to chromatin relaxation (**B**) Chromatin relaxation in U2OS *WT*, *PARP1*^*KO*^, *HPF1*^*KO*^ and *PARP1/HPF1* double knockout cells, assessed by the thickness of the highlighted damaged chromatin line at 120 s relative to 0 s post irradiation. (**C**) Chromatin relaxation at 120 s post irradiation in *WT* and *PARP1*^*KO*^ cells expressing mCherry-PARP1 WT or PARP1-LW/AA. ⵁ denotes no plasmid expression. (**D**) Chromatin relaxation at 120 s post-irradiation in *WT* and *HPF1*^*KO*^ cells expressing mCherry-HPF1 D283A. ⵁ denotes no plasmid expression. (**E**) Kinetics of chromatin relaxation in U2OS *WT* cells overexpressing or not mCherry-HPF1. Data are shown as mean ± SEM.

Interestingly, we also found that the overexpression of wild-type HPF1 led to a dramatic increase of the chromatin relaxation (Fig 3E). This massive unfolding of the chromatin structure remains nevertheless reversible. Indeed, a progressive recondensation of the chromatin structure was also observed in HPF1 overexpressing cells following the initial relaxation phase albeit slower than for endogenous levels of HPF1. Approximately 30 min post irradiation, the chromatin compaction levels were similar to those pre-damage in these cells. Altogether, the findings presented in this section reveal that HPF1 plays a central role in the PARP1-dependent chromatin remodeling events occurring at early stages of the DDR.

### Chromatin remodeling triggered by PARP1/HPF1 at DNA lesions does not depend on auto APD-ribosylation of PARP1 but rather relies on trans ADP-ribosylation of histones

To further investigate the mechanisms underlying the regulation of chromatin remodeling by HPF1 at sites of DNA damage, we first analyzed the behaviour of the HPF1 mutant E284A, which still interacts with PARP1 but blocks the ADP-ribosylation activity of the joined catalytic site created by the PARP1/HPF1 heterodimer^9^. While this E284A mutant displayed increased recruitment to DNA lesions compared to wild-type HPF1 (Fig S3B) in line with its tighter binding to PARP1^12^, it was unable to rescue the chromatin relaxation defect observed in *HPF1*^*KO*^ cells (Fig 4A). This finding demonstrates that the ADP-ribosylation activity of the PARP1/HPF1 complex is needed for chromatin remodeling at sites of damage.

**Figure 4:**
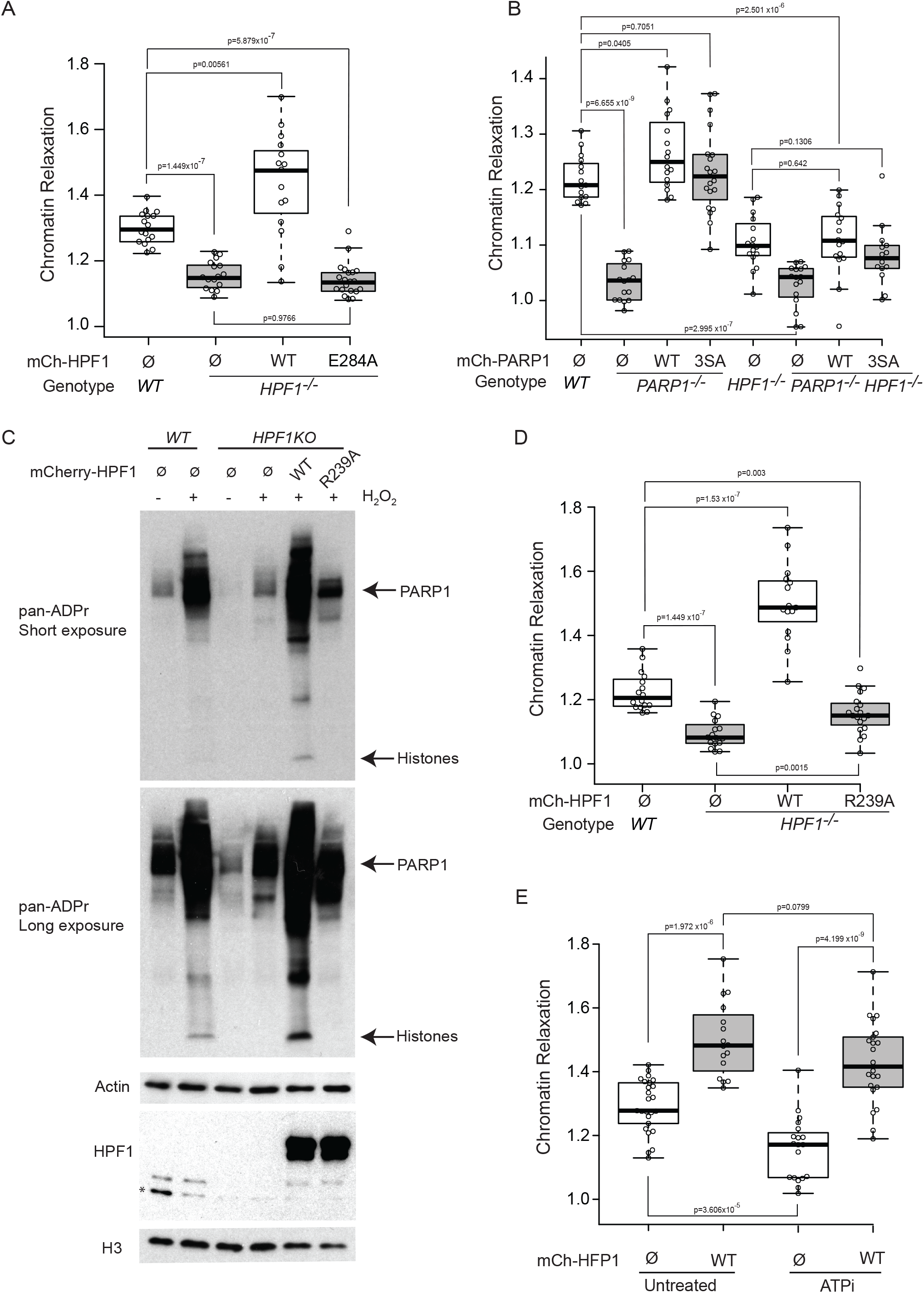
HPF1-dependent chromatin relaxation relies on trans ADP-ribosylation of histones rather than PARP1 auto-modification. (**A**) Chromatin relaxation at 120 s post-irradiation in *WT* and *HPF1*^*KO*^ cells expressing or not mCherry-HPF1 E284A. ⵁ denotes no plasmid expression. (**B**) Chromatin relaxation at 120s post-irradiation in U2OS *WT*, *PARP1*^*KO*^, *HPF1*^*KO*^ and *PARP1/HPF1* double knockout cell. Cells are complemented either with mCherry-PARP1 WT or with PARP1-3SA mutant. ⵁ denotes no plasmid expression. (**C**) Western blot displaying ADPr signals, stained with a pan-ADPr antibody, in *WT* and *HPF1*^*KO*^ cells expressing HPF1-WT or HPF1-R239A and treated or not with H_2_O_2_. H3 and Tubulin were used as loading controls. (**D**) Chromatin relaxation at 120 s post-irradiation in *WT* and *HPF1*^*KO*^ cells expressing mCherry-HPF1 R239A. ⵁ denotes no plasmid expression. (**E**) Chromatin relaxation at 120 s post-irradiation in U2OS *WT* cells overexpressing mCherry-HPF1 and depleted or not for ATP (ATPi). ⵁ denotes no plasmid expression.

Upon DNA damage, HPF1 has been shown to control the addition of ADP-ribose moieties on the serine residues of specific targets^10^. This includes the auto ADP-ribosylation of PARP1 itself but also trans ADP-ribosylation of other targets, in particular histones which are the main ADP-ribosylation targets after PARP1^10^. Therefore, we assessed the relative contributions of the ADP-ribosylation of PARP1 and the histones to the HPF1-dependent chromatin relaxation observed at sites of damage. First, we found that the PARP1-3SA mutant, which can still catalyze histone ADP-ribosylation due to interaction with HPF1^16^ but shows greatly reduced auto ADP-ribosylation (Fig S1B,C), restored chromatin relaxation in *PARP1*^*KO*^ cells similar to PARP1-WT complementation (Fig 4B). As expected, this rescue relied on the presence of HPF1 since expressing PARP1-3SA in *PARP1*^*KO*^/*HPF1*^*KO*^ cells did not permit chromatin relaxation to reach the level observed in WT cells. Next, we analysed the behaviour of the HPF1 mutant R239A at DNA lesions. In agreement with *in vitro* observations showing that this mutation does not significantly impact the interaction with PARP1^13^, we observed that HPF1-R239A was recruited to DNA lesions, although at a lower level than its wild-type counterpart (Fig S4A,B). Furthermore, this mutation perturbed the catalytic activity of the PARP1/HPF1 complex by preventing histone ADP-ribosylation while preserving some PARP1 automodification (Fig 4C), confirming previous reports^9,13^. When expressed in *HPF1*^*KO*^ cells, the HPF1-R239A mutant was unable to promote chromatin relaxation as observed with wild-type HPF1 complementation (Fig 4D). Together with the results regarding the PARP1-3SA mutant, these findings indicate that the driving force for chromatin remodelling at sites of damage is the ADP-ribosylation of histones rather than of PARP1.

Finally, given that several chromatin remodelers were found to recruit to DNA lesions and contribute to chromatin remodeling at sites of damage^23–25^, we wondered whether the chromatin relaxation promoted by histone ADP-ribosylation relied on ATP-dependent processes. ATP-deprivation, which leaves ADP-ribosylation signalling unaffected at sites of damage^23^, did not prevent the dramatic increase in chromatin relaxation observed upon overexpression of HPF1 (Fig 4E). Therefore, histone ADPr appears sufficient to promote chromatin unfolding, without the need for active remodeling processes.

### HPF1-dependent chromatin relaxation at DNA lesions promotes the recruitment of repair factors

Previous studies have indicated that ADPr-dependent chromatin remodeling contributes to the efficient recruitment of several repair factors through increased accessibility to DNA at sites of damage^24–26^. Given the central role of HPF1 in these chromatin remodeling events, we investigated the impact of the loss of HPF1 on the recruitment of two repair factors, CHD4 and CHD7, that were both reported to accumulate at DNA lesions in response to ADPr-dependent chromatin relaxation. Importantly, these two factors belong to different repair pathways with CHD4 involved in homologous recombination (HR)^27^ while CHD7 participates in non-homologous end-joining (NHEJ)^24^. The accumulation of these two factors to sites of damage was impaired in both *PARP1*^*KO*^ and *HPF1*^*KO*^ cells, the strength of the recruitment defect mirroring the reduction in chromatin relaxation observed in both cell lines (Fig 5A-D). Furthermore, expressing PARP1-3SA in *PARP1*^*KO*^ cells rescued the accumulation of CHD4 and CHD7 to sites of damage similar to PARP1-WT complementation in contrast to the expression PARP1-LW/AA in *PARP1*^*KO*^ or to the complementation of *PARP1^KO^/HPF1^KO^* cells with PARP1-WT (Fig S5A-D). These findings show that the recruitment defect observed for CHD4 and CHD7 in *HPF1*^*KO*^ cells is not the consequence of impaired automodification of PARP1 but is rather due the reduced chromatin relaxation associated with the absence of HPF1-dependent histone ADP-ribosylation at sites of damage.

**Figure 5:**
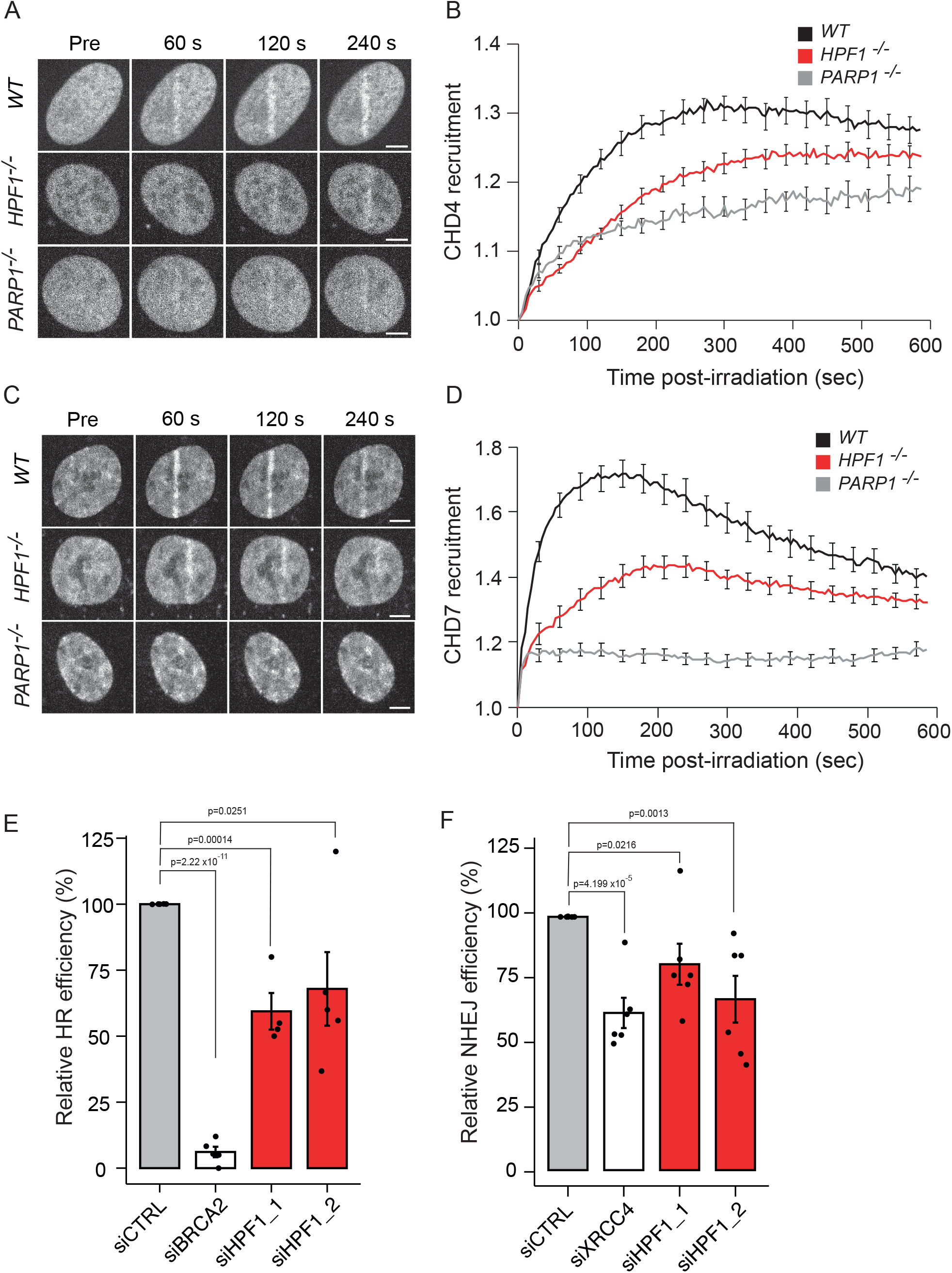
HPF1-dependent chromatin relaxation promotes the recruitment of both homologous recombination and non-homologous end joining repair factors. (**A**) Representative confocal images showing recruitment of GFP-CHD4 to sites DNA damage induced by laser irradiation, in U2OS *WT*, *HPF1*^*KO*^ or *PARP1*^*KO*^ cells. Scale bar, 5 μm. (**B**) Recruitment kinetics of GFP-CHD4 to sites of DNA damage in U2OS *WT*, *HPF1*^*KO*^ or *PARP1*^*KO*^ cells. (**C**) Representative confocal images showing recruitment of GFP-CHD7 to sites of DNA damage inducted by laser irradiation, in U2OS *WT*, *HPF1*^*KO*^ or *PARP1*^*KO*^ cells. Scale bar, 5 μm. (**D**) Recruitment kinetics of GFP-CHD7 to sites of DNA damage in U2OS *WT*, *HPF1*^*KO*^ or *PARP1*^*KO*^ cells. (**E**) Quantification of DR-GFP-positive U2OS cells transfected with the indicated siRNA and I-SceI expression vector. The mean ± SEM of 5 independent experiments is shown. Data were normalized to siCTRL, which was set to 100%. (**F**) Quantification of EJ5-GFP-positive U2OS cells transfected with the indicated siRNA and I-SceI expression vector. The mean ± SEM of 6 independent experiments is shown. Data were normalized to siCTRL, which was set to 100%.

Previous data has shown that HPF1-deficient cells are hypersensitive to DNA damaging agents, suggesting that DNA repair efficiency is compromised^8^. To better characterize this repair defect, we depleted HPF1 in the well-established reporter cell lines DR-GFP and EJ5-GFP and found that reducing HPF1 levels impaired the efficiency of DNA repair by both HR and NHEJ (Fig. 5E,F and Fig. S5E-H). This result is in line with the fact that repair factors involved in each of these pathways displayed reduced recruitment at DNA lesions in *HPF1*^*KO*^ cells (Fig 5A-D).

## DISCUSSION

ADP-ribosylation is one of the earliest signalling pathways activated during the DDR^28^. It is well-established that PARP1 is the central engine triggering this signalling pathway via its rapid recruitment to DNA lesions^2^ but recent reports have demonstrated that this process also requires a steering wheel, the co-factor HPF1, to dictate the choice of the target proteins that will be ADP-ribosylated^9–11^. HPF1 was shown to associate with PARP1 to form a joint catalytic site that is essential to ADP-ribosylate serine residues of target proteins such as PARP1 itself or histones^9^. Nevertheless, the fact that HPF1 is much less abundant than PARP1 within the cell nucleus suggests that the regulatory role played by HPF1 does not require a stable association between the two partners but is rather exerted via transient interaction with PARP1, which is needed to attach the first ADPr moiety on the serine residues^9,15,16^. Yet, the factors regulating this temporary association between PARP1 and HPF1, and therefore, controlling HPF1 accumulation to sites of damage, remain only partially described. In this report, we demonstrate that the rapid accumulation of HPF1 to sites of damage fully relies on its binding to PARP1 since impairing this interaction by mutating either the last two amino acids of PARP1 or the residue D283 of HPF1 suppressed HPF1 recruitment to sites of damage (Fig 1, Fig S3B). We also found that, while HPF1 mobilization from sites of damage is influenced by the release of PARP1 (Fig 1F,G), the dissipation speed of HPF1 is slower than that of PARP1 (Fig 1C) leading to a gradual increase of the HPF1/PARP1 ratio at sites of damage. This finding hints for progressive changes in the characteristics of ADP-ribosylation signalling along the course of the DDR. While the early wave of ADP-ribosylation could be mainly composed of HPF1-independent PARP1 auto-modification, trans-ADP-ribosylation of histones triggered by PARP1/HPF1 would arise at later stages of the DDR. Besides its role to target the residues for ADP-ribosylation, HPF1 also controls the length of the ADP-ribose chains^12^. Here, we show that these *in vitro* data hold true in living cells and that the accumulation of HPF1 at sites of damage favours the addition of ADPr chains but restricts their length (Fig 2). This damping function of HPF1 is likely a central regulator of ADP-ribosylation signalling given the prevalence of mono-ADPr modifications compared to poly-ADPr chains in the DDR context^11^.

The ADPr moieties bound to target proteins located nearby the DNA lesions are recognized by multiple effectors contributing to the restoration of genomic integrity^29^. While this role as a binding platform has been studied extensively^29^, less is known about the direct impact of these ADPr chains on the function of the proteins to which they are attached to. More specifically, while auto-modification of PARP1 has been shown to be important for the timely mobilization of this protein from sites of damage^16,30^, the direct impact of trans-ADP-ribosylation, in particular, on histones, has not been elucidated. In this current report, we demonstrate that HPF1-dependent histone ADP-ribosylation is a major contributor to the transient unfolding of the chromatin in the vicinity of the DNA lesions (Fig 3,4). Seminal *in vitro* work by Poirier and colleagues had found that the ADP-ribosylation of histones was sufficient to decondense purified chromatin fibers^31^, a process that does not require histone eviction ^32,33^. Our live-cell results are fully consistent with these data. Indeed, we previously found that ADPr-dependent chromatin relaxation that occurs at early stages of the DDR does not involve the mobilization of core histones^22^. Additionally, we now show that HPF1-dependent histone ADP-ribosylation promotes chromatin unfolding even upon ATP-deprivation, suggesting that the addition of the ADPr onto histones themselves is sufficient to promote unfolding without the need for active nucleosome disassembly via chromatin remodelers (Fig 4E). Also in agreement with *in vitro* results^33^, the dynamic nature of ADPr, which can be removed by different hydrolases such as PARG or ARH3^29,34,35^, allows the chromatin relaxation process to be fully reversible even in the context of HPF1 overexpression which strongly enhances this relaxation (Fig 3E). Importantly, several ATP-dependent remodelers have also been shown to contribute to early chromatin relaxation at sites of damage^23–25,36^. While future work will help to define whether these two different modalities of chromatin remodeling are coordinated or work independently, it was recently shown that histone ADP-ribosylation can serve as a potential docking site for some remodelers such as ACL1/CHD1L^37^ known to contribute to chromatin remodeling at sites of damage^23^.

We recently proposed that the PARP-dependent chromatin relaxation observed at early stages of the DDR facilitates access of DNA lesions to repair factors^22^. In the present work, we demonstrate that this process is triggered by trans-ADP-ribosylation of histones rather than auto-modification of PARP1 and that it promotes the accumulation of repair factors belonging to both HR and NHEJ. The fact that HPF1 deficiency impairs the efficient of both repair pathways indicates that these chromatin remodelling processes regulated by the PARP1/HPF1 axis precedes the repair pathway choice and therefore are pivotal in the initiation of the DDR (Fig 6).

**Figure 6:**
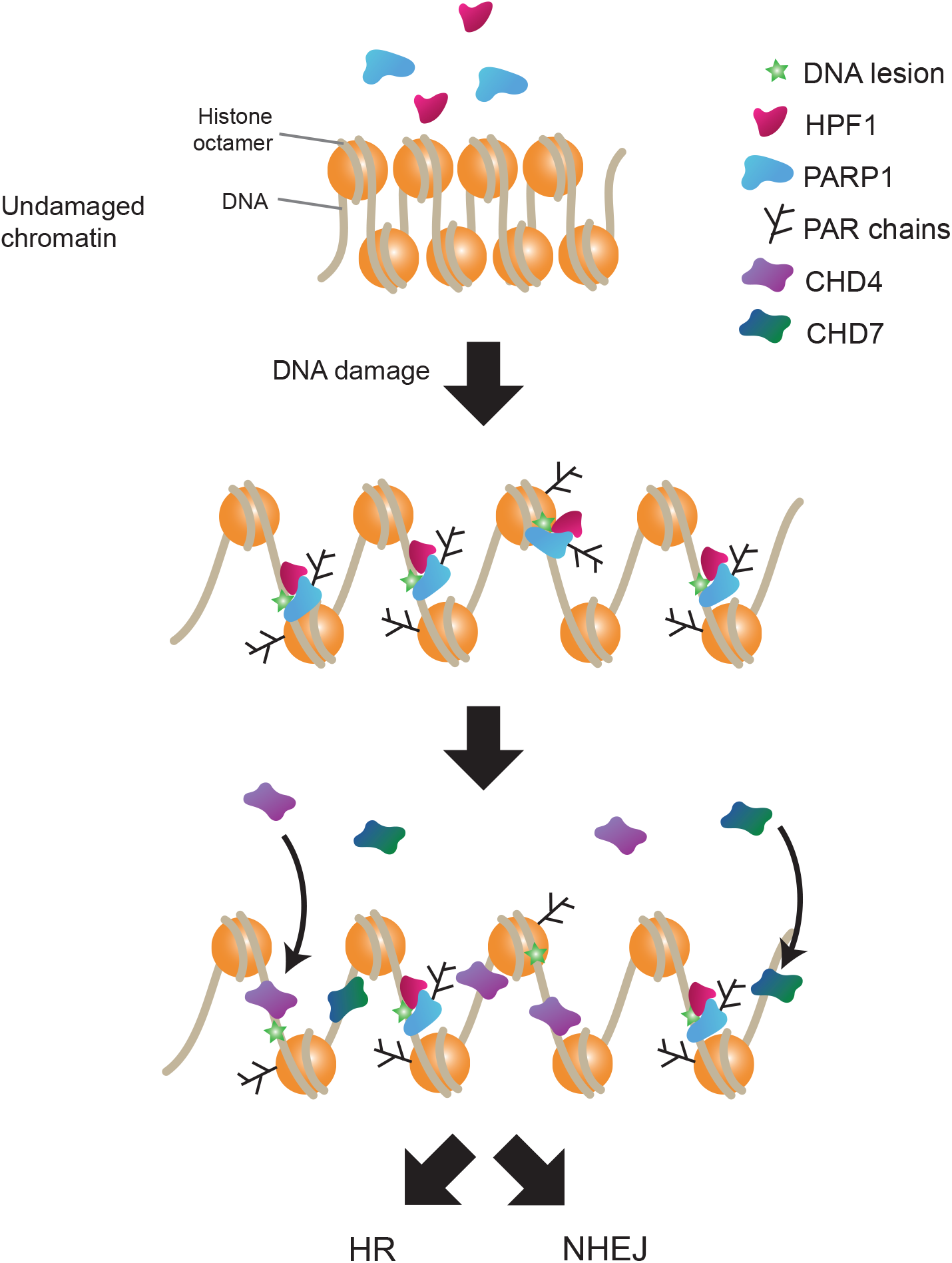
HPF1 regulates DNA damage induced chromatin relaxation. Upon damage, HPF1 interacts with PARP1 at sites of damage and promote both PARP1 auto-modification and trans ADP-ribosylation of histone. Histone ADP-ribosylation promotes chromatin relaxation in the vicinity of DNA lesions, promoting the recruitment of repair factors to facilitate genome restoration by both homologous recombination and non-homologous end joining.

## Acknowledgements

We thank the Microscopy-Rennes Imaging Center (BIOSIT, Université Rennes 1), member of the national infrastructure France-BioImaging supported by the French National Research Agency (ANR-10-INBS-04), for providing access to the imaging setups, as well as S. Dutertre and X. Pinson for technical assistance on the microscopes. We also thank the Cytométrie en flux et tri cellulaire (BIOSIT, Université Rennes 1), specifically L. Deleurme and A. Amie for technical assistance with flow cytometry. We would also like to thank I. Ahel and M. Suskiewicz for thoughtful discussions and generously sharing HPF1 plasmid DNA. For this work, the S.H.’s group received financial support from the Agence Nationale de la Recherche (PRC-2018 REPAIRCHROM), the Institut National du Cancer (PLBIO-2019) and the Institut Universitaire de France, to S.H. R.S. is supported by the Fondation ARC pour la recherche sur le cancer (PDF20181208405). The work in the Timinszky laboratory was supported by the Hungarian Academy of Sciences (LP2017-11/2017) and the National Research Development and Innovation Office (K128239).

## Author Contributions

R.S., G.T. and S.H. conceived the project with inputs from all authors. R.S., S.Z. and S.H. performed live-cell imaging and analysed the imaging data. S.Z. performed the DNA repair assay. C.C. and R.S. generated DNA constructs. R.S., G.T. and S.H. wrote the paper with input from all authors.

## Methods

### Plasmids

pmEGFP-WWE, PATagRFP-H2B^23^, pH2B-PAGFP^38^, pGFP-CHD4^39^, pGFP-CHD7^24^, pPARP1-mCherry^20^, pmCherry-PARP1 WT, pmCherry-PARP1 E988K^17^, pLacI-GFP trap^25^, pcDNA3.1(+) (Thermofisher) and pmCherry-C1 (Takara) were previously described. pCBASceI was a gift from Maria Jasin (Addgene plasmid # 26477,^40^). PARP1 3SA (S499A/S507A/S519A) cDNA was amplified from pDEST-YFP-PARP1-3SA^16^ and ligated into pmCherry-C1 between *Bgl*II and *Xma*I. pmCherry-PARP1 L1013A/W1014A was made using site-directed mutagenesis with primers provided in Table 1. MacroH2A1.1 macrodomain cDNA was amplified from pcDNA3.1-YFP-macroH2A1.1 macrodomain^20^ and ligated into pEGFP-C1 between *Bgl*II and *Eco*RI. cDNA of HPF1 WT, R239A, D283A and E284 were amplified from pDEST-YFP-HPF1 WT, R239A, D283A and E284^9^ with primers provided in Table 1 and ligated into pEGFP-C1 or pmCherry-C1 between *Bgl*II and *Bam*HI.

**Table 1:**
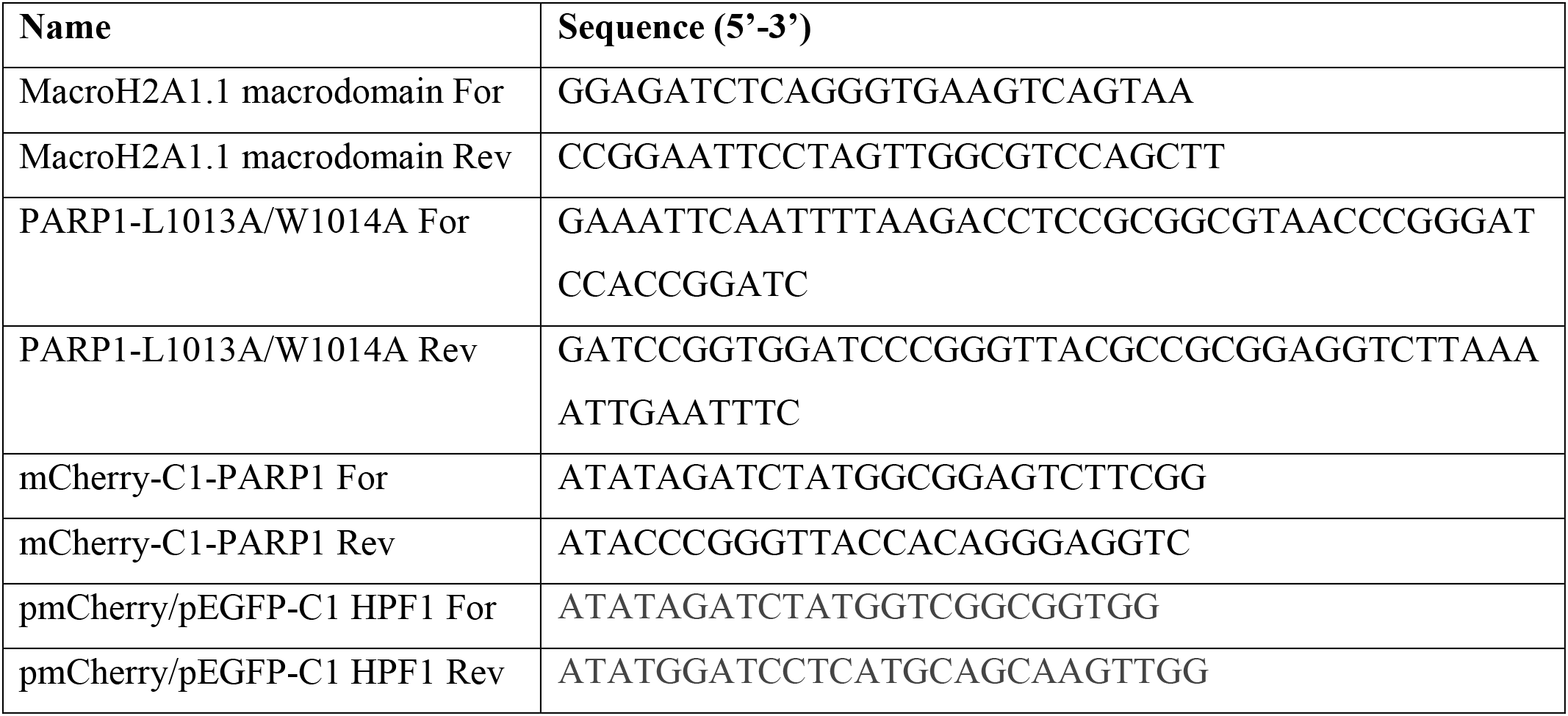
List of primers used in this study

### Cell culture

All cells used in this study were cultured in DMEM (Sigma) or RPMI supplemented with 10% FBS, 100 μg/mL penicillin, 100 U/mL streptomycin and maintained at 37°C in a 5% CO_2_ incubator. U2OS *WT*, U2OS *PARP1*^*KO*^, U2OS *HPF1*^*KO*^, U2OS *PARP1^KO^ HPF1^KO^* double knockout cells were generated previously^8^. U2OS 2B2 were generated previously^41^. U2OS-DR and U2OS-EJ5 cells were a kind gift from Jeremy Stark^42^. All experiments presented in this work were performed on unsynchronized cells.

### Live-cell microscopy

U2OS cells were seeded into 8-well Imaging Chamber CG (Zell-Kontakt) and transfected 48-72 h prior to imaging using XtremeGene HP (Roche) according to the manufacture’s instructions. Cells were sensitized by aspirating growth medium from the Lab-Tek and replacing it with fresh medium containing 0.15 – 0.3 μg/mL Hoechst 33342 for 1 hour at 37°C. Immediately prior to imaging, growth medium was replaced with CO_2_-independent imaging medium (Phenol Red-free Leibovitz’s L-15 medium (Life Technologies) supplemented with 20% fetal bovine serum, 2 mM glutamine, 100 µg/mL penicillin and 100 U/mL streptomycin). ATP depletion was achieved by bathing the cells for at least 30 min with PBS containing 10% fetal bovine serum, 10 mM NaN3 and 50 mM 2-deoxyglucose (Platani et al., 2002). Live-cell imaging experiments were completed on a Ti-E inverted microscope from Nikon equipped with a CSU-X1 spinning-disk head from Yokogawa, a Plan APO 60x/1.4 N.A. oil-immersion objective lens and a sCMOS ORCA Flash 4.0 camera. The fluorescence of EGFP/PAGFP and mCherry/PATagRFP were excited with lasers at 490 and 561 nm, respectively. For fluorescence detection, we used bandpass filters adapted to the fluorophore emission spectra. Laser microirradiation and local photoactivation at 405 nm was performed along a 16 μm-line through the nucleus using a single-point scanning head (iLas2 from Roper Scientific) coupled to the epifluorescence backboard of the microscope. To ensure reproducibility laser power at 405 nm was measured at the beginning of each experiment and set to 125 μW at the sample level. Cells were maintained at 37°C with a heating chamber. Protein recruitment was quantified using a custom-made MATLAB (MathWorks) routine which measures the mean intensity within the damaged region (*I*_*d*_) as determined by the segmentation of the photoactivated H2B signal, the mean nuclear fluorescence (*I*_*nd*_), and the mean background signal outside of the cell (*I*_*bg*_). Protein accumulation at sites of damage (*A*_*d*_) was then calculated as:

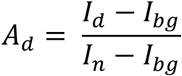

The intensity within the microirradiated area was then normalized to the intensity prior to damage induction.

Chromatin relaxation was determined using a custom MATLAB routine that measures the changes in the thickness of the photoconverted H2B line relative to its value immediately after damage induction^23^.

The PAR3H assay has been previously described^25^. Briefly, U2OS-2B2^41^ cells containing the LacO array were transfected with GFP-macrodomain of macroH2A1.1, LacI-GFP trap and mCherry-PARP1, mCherry-PARP1 E988K or mCherry-PARP1 3SA. Cells sensitized with Hoechst 33342 were irradiated away from the LacO array with 405 nm light to induce DNA damage as described above. No matter whether it is ADP-ribosylation status, PARP1 does not remain stably bound to DNA lesions but can quickly dissociate from this region and diffuse within the nucleus^17,18^. Therefore, the amount of accumulation of mCherry tagged PARP1 at the LacO array due to interaction with tethered macrodomain can then be used as a proxi to assess the level of ADP-ribosylation of the different PARP1 mutants. The mCherry intensity at the LacO array (*A*_*lo*_) quantified with the following equation where *I*_*o*_ is the intensity of the LacO array, *I*_*n*_ is the mCherry signal in the nucleoplasm devoid of the LacO array and *I*_*bg*_ is the intensity of the background:

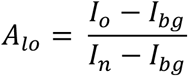

The intensity within the LacO array was then normalized to the intensity prior to damage induction.

### DNA repair assay

U2OS-DR and U2OS-EJ5 cells containing either a stably integrated cassette of the DR-GFP or EJ5-GFP reporter were used to measure the repair of I-SceI-induced DSBs by homologous recombination or by non-homologous end joining, respectively^42^. Briefly, cells were transfected with siRNA for 48 h prior to co-transfected with an mCherry expression vector and the I-SceI expression vector. The percentage of GFP-positive cells among the mCherry-positive cells was determined 48 h after I-SceI transfection using a LSRFortessa X-20 (BD Bioscience) using BD FACSDiva Software v8.0.1. Quantifications were performed with FACSDiva™ (BD Biosciences). siRNAs used in this study are shown in Table 2.

**Table 2:**
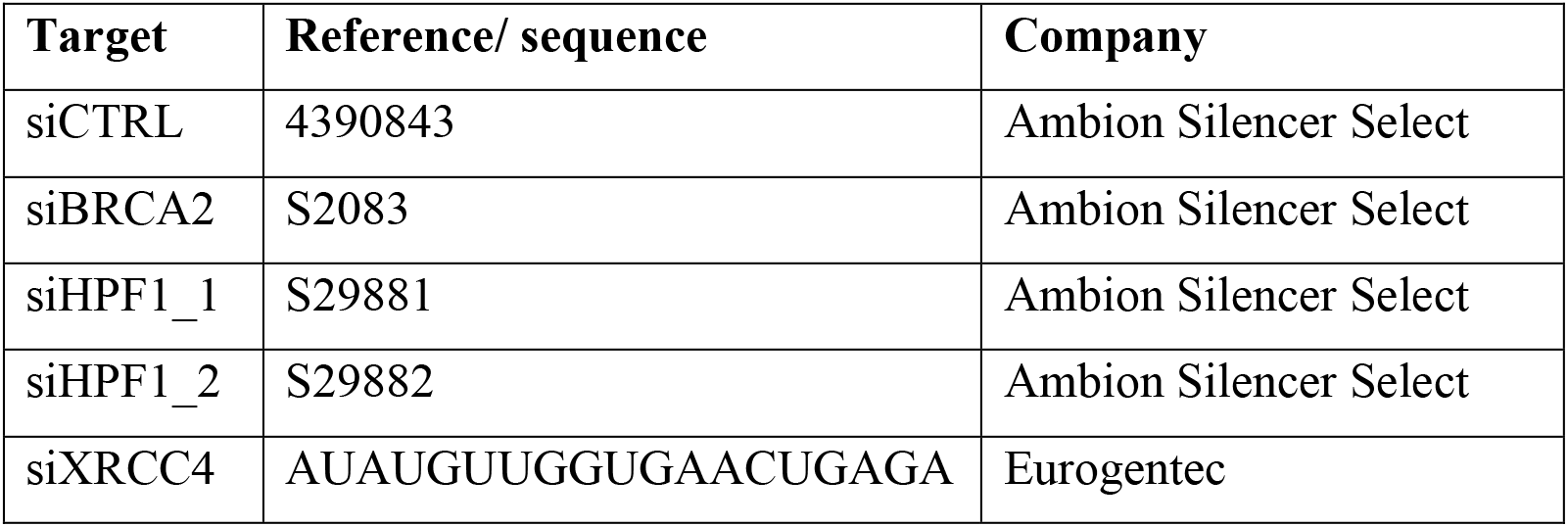
List of siRNA used in this study

### Western Blotting

For whole cell extract, cells were lysed on with Triton-X buffer (% Triton X-100, 100 mM NaCl and 50 mM Tris-HCl, pH 8.0, 5 mM MgCl_2_, 0.1% Benzonase (Sigma Aldrich), 1x protease inhibitor (Roche) on an orbital rotator at 4°C for 30 min. Samples were centrifuged at 20 000 g for 15 min and supernatant was collected. Protein samples were quantified using Bradford (BioRad) and equal amounts of protein were loaded on gels for SDS-PAGE prior to immunoblotting. Antibodies used in this study are given in Table 3. For DNA damage induction, cells were incubated in serum free media prior to treatment with 2 mM H_2_O_2_ for 10 min. During cell lysis, Triton-X buffer was supplemented further with 2 μM olaparib (Selleckchem) and 2 μM PARG inhibitor PDD00017273 (Sigma Aldrich)

**Table 3:** List of antibodies used in this study

| Target | Host | Company | Reference | Dilution in WB |
| --- | --- | --- | --- | --- |
| **Primary Antibodies** |  |  |  |  |
| PARP1 | Rabbit | - | Sellou et al. <sup>23</sup> | 1:10000 |
| PARP2 | Rabbit | Active Motif | #39744 | 1:1000 |
| HPF1 | Rabbit | Novus<br>Biologicals | NBP1-93973 | 1:500 |
| Actin | Mouse | Sigma | A5060 | 1:1000 |
| XRCC4 | Mouse | Abcam | Ab213729 | 1:500 |
| BRCA2 | Rabbit | Novus<br>Biologicals | MAB2476 | 1:500 |
| RFP/mCherry | Mouse | Chromotek | 6g6-100 | 1:1000 |
| H3 | Rabbit | Abcam | Ab1731 | 1:2500 |
| PAN- ADPr binding reagent | Rabbit | Sigma | MABE1016 | 1:1500 |
| **Secondary Antibodies** |  |  |  |  |
| Anti-Mouse-HRP | Goat | Agilent | P044701-2 | 1:3000 |
| Anti-Rabbit-HRP | Swine | Agilent | P039901-2 | 1:3000-1:8000 |

### Statistics

Data analysis and visualization was performed using R software (https://www.r-project.org/). The boxplot limits correspond to the 25^th^ and 75^th^ percentiles and the bold line indicates the median value. The whiskers extend 1.5 times the interquartile range. The timelapse curves are the mean ± SEM of at least 15 cells per condition from a characteristic experiment among at least three independent repeats. The histograms show the mean ± SEM of the indicated independent experiments number in the figure legends. Unless stated otherwise, p values were calculated using an unpaired Student’s t test, assuming unequal variances.

## Supplementary figure legends

**Supp Figure 1:**
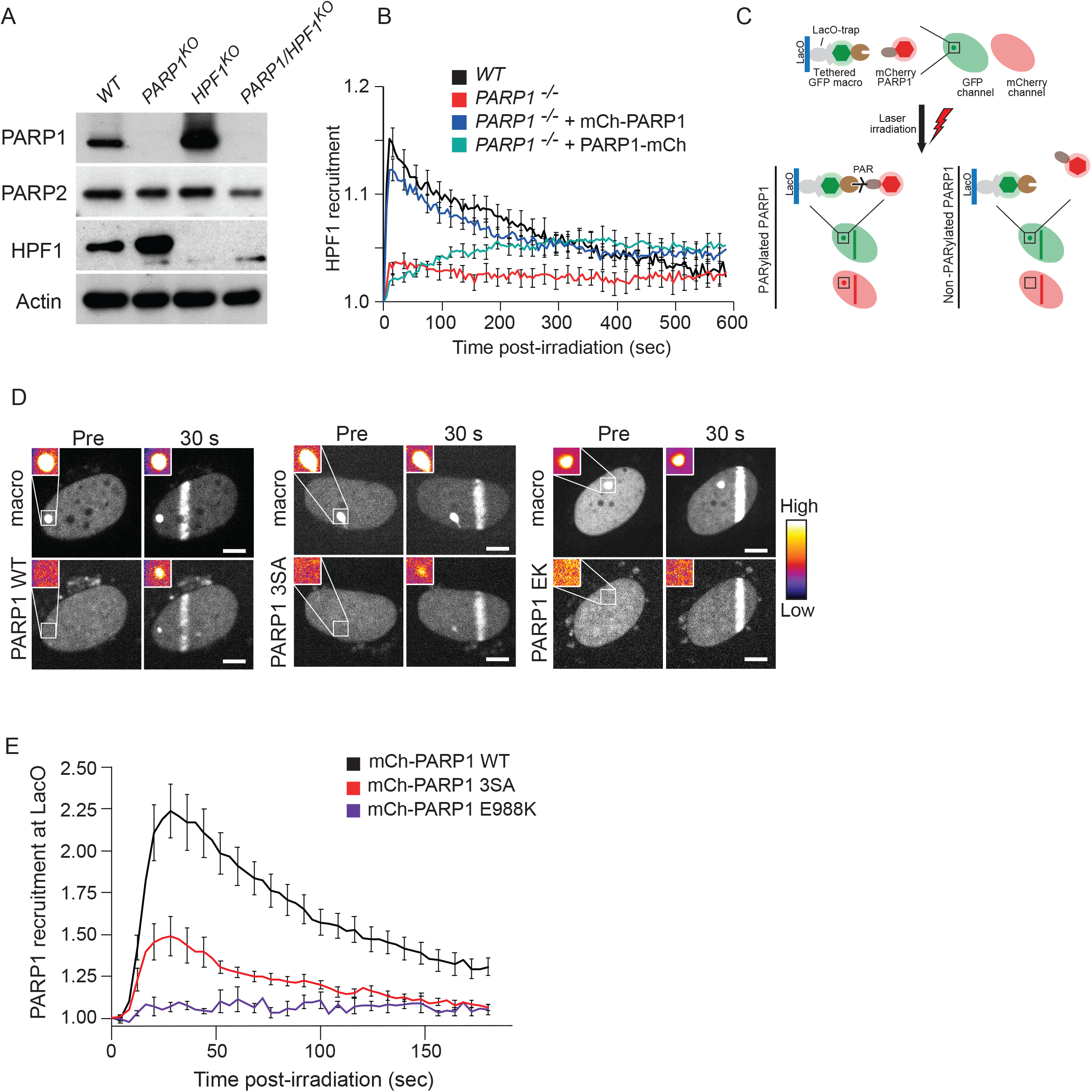
HPF1 recruitment to sites of damage relies on interaction with the C-terminus of PARP1. (**A**) Immunoblots of whole cell extract from U2OS *WT*, *PARP1*^*KO*^, *HPF1*^*KO*^ and *PARP1/HPF1* double knockout cells. Actin is used as a loading control. (**B**) Recruitment kinetics of GFP-HPF1 to sites of DNA damage in *WT* or *PARP1*^*KO*^ cells expressing WT N-terminally (mCh-PARP1) and C-terminally tagged PARP1 (PARP1-mCh). Data are shown as mean ± SEM. (**C**) Schematic representation of PAR-3H assay. In this assay, mCherry-tagged PARP1 variants are expressed together with the GFP-tagged macrodomain of macroH2A1.1, that is tethered to a LacO array integrated into the genome of U2OS 2B2 cells. Upon laser irradiation, the PARP1 variants recruit to sites of damage, where they can be auto ADP-ribosylated or not, depending on the variant. A defect in auto ADP-ribosylation does not preclude high turnover at sites of damage, allowing all PARP1 variants to diffuse rapidly within the nucleus after their release from the DNA lesions. If PARP1 is ADP-ribosylated it can then interact with the tethered macrodomain leading to an increase in mCherry signal at the LacO array. (**D**) Representative confocal images of PARP1-WT, PARP1-3SA or PARP1-E988K to YFP-macrodomain of mH2A1.1 tethered to LacO. Inset, pseudocolored according to the look-up table displayed, shows the magnified LacO array. Post-irradiation images are shown at 30 seconds. Scale bar, 5 μm. (**E**) Quantification of the accumulation of PARP1-WT, PARP1-3SA or PARP1-E988K to the LacO array after DNA induction by laser irradiation.

**Supp Figure 2:**
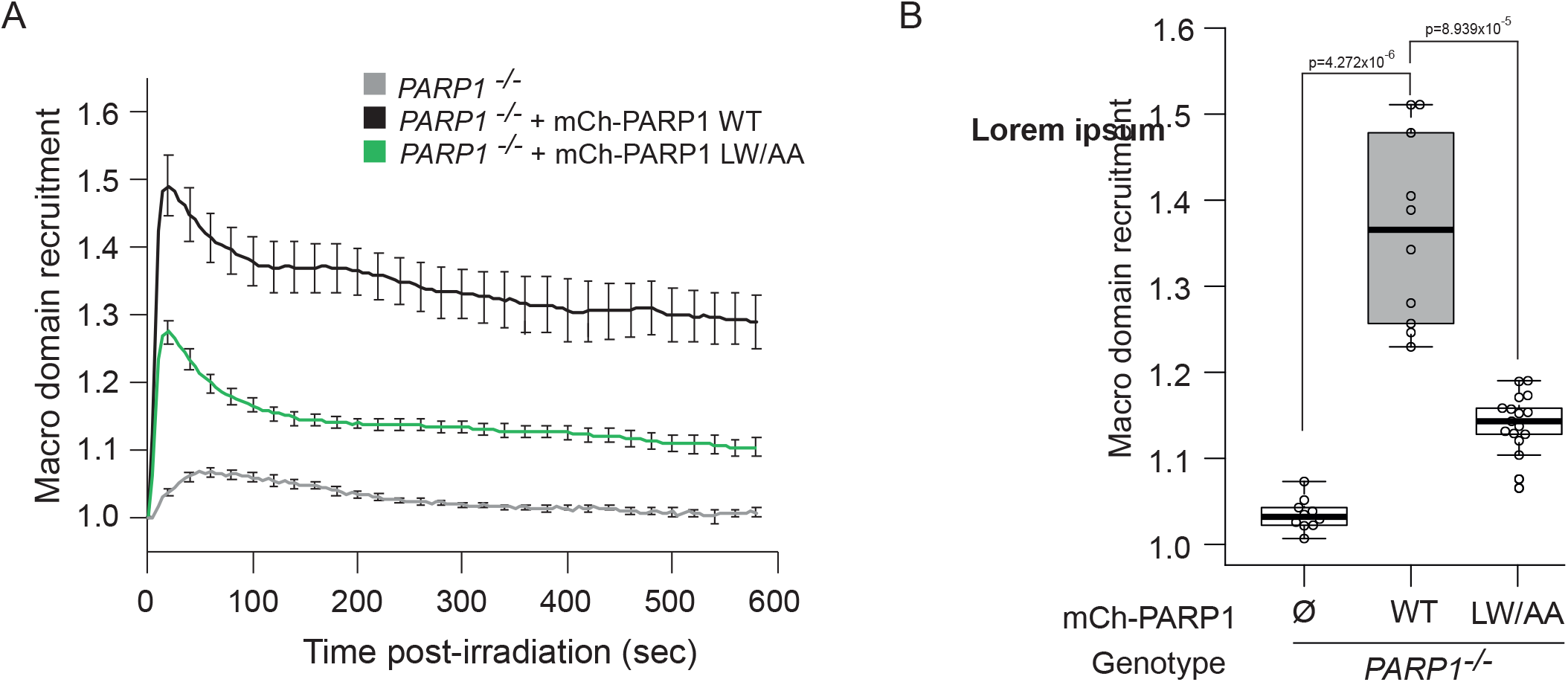
HPF1 regulates ADP-ribosylation signalling at sites of DNA damage. **(A)** Recruitment kinetics of GFP-macrodomain of mH2A1.1 at sites of DNA damage induced by laser irradiation, in U2OS *PARP1*^*KO*^ cells complemented or not with mCherry-PARP1 WT, PARP1-3SA or PARP1-LW/AA. (**B**) Quantification of mean intensity of GFP-macrodomain of mH2A1.1 at sites of DNA damage 200 s post-irradiation in *PARP1*^*KO*^ complemented or not with mCherry-PARP1 WT or PARP1-LW/AA mutants. ⵁ denotes no plasmid expression.

**Supp Figure 3:**
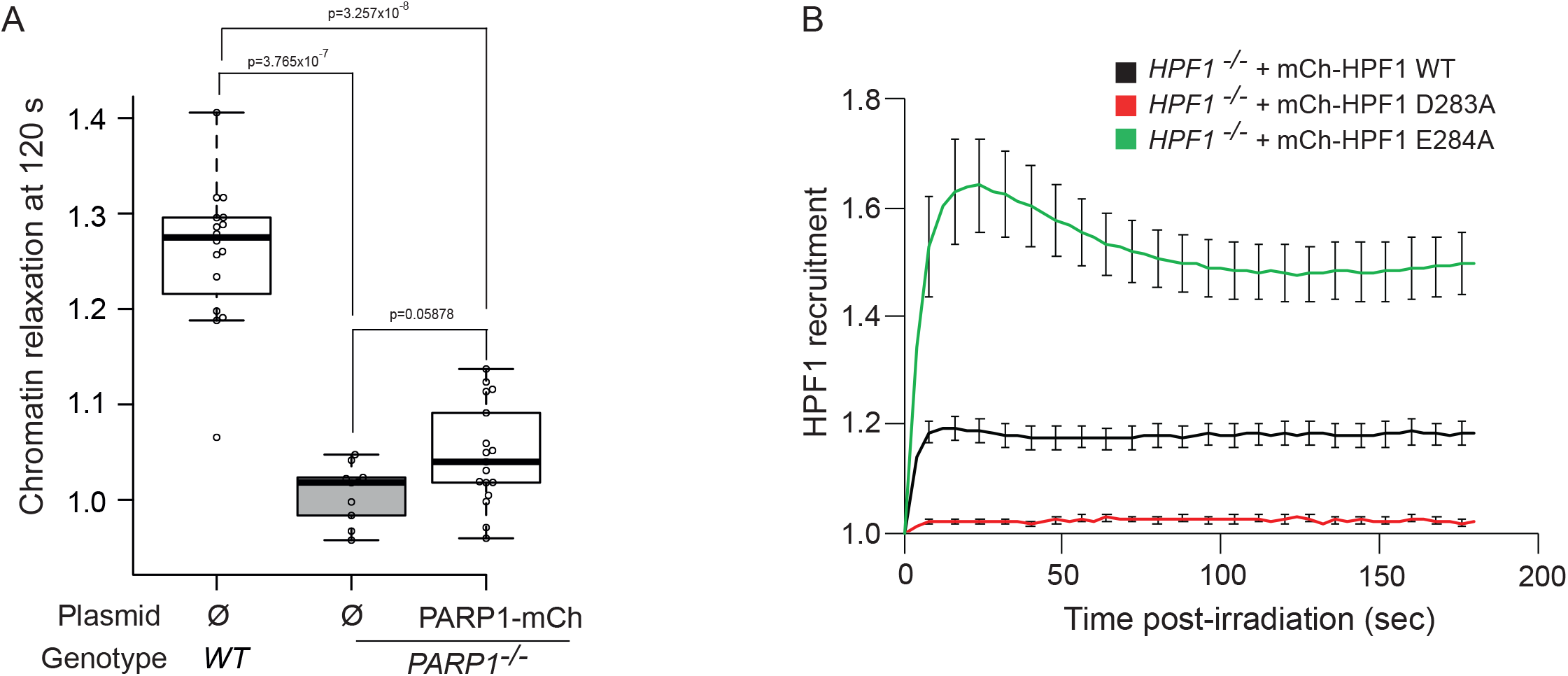
HPF1 promotes chromatin relaxation at sites of DNA damage. (**A**) Chromatin relaxation in U2OS *WT* or *PARP1*^*KO*^ cells at 120 s post-irradiation. Cells are complemented or not with C-terminally-tagged PARP1-mCherry. ⵁ denotes no plasmid expression. (**B**) Recruitment kinetics of mCherry-tagged HPF1 WT and the point mutants D283A and E284A at sites of DNA damage in U2OS *HPF1*^*KO*^ cells.

**Supp Figure 4:**
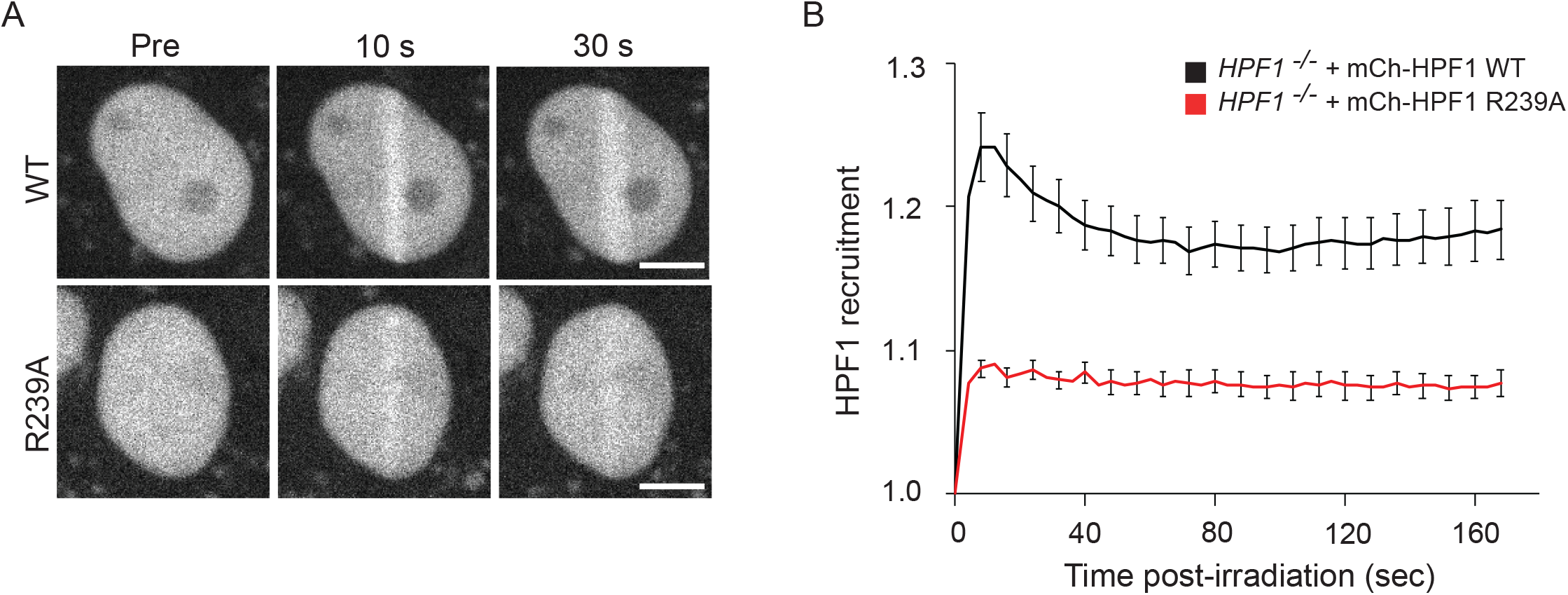
HPF1-dependent chromatin relaxation relies on trans ADP-ribosylation of histones rather than PARP1 auto-modification. **(A)** Representative images of the recruitment of mCherry-tagged HPF1 WT or HPF1-R239A to sites DNA damage induced by laser irradiation in U2OS *HPF1*^*KO*^ cells. Scale bar, 5 μm. (**B**) Recruitment kinetics of mCherry-tagged HPF1 WT or HPF1-R239A mutant at sites of DNA damage in U2OS *HPF1*^*KO*^ cells.

**Supp Figure 5:**
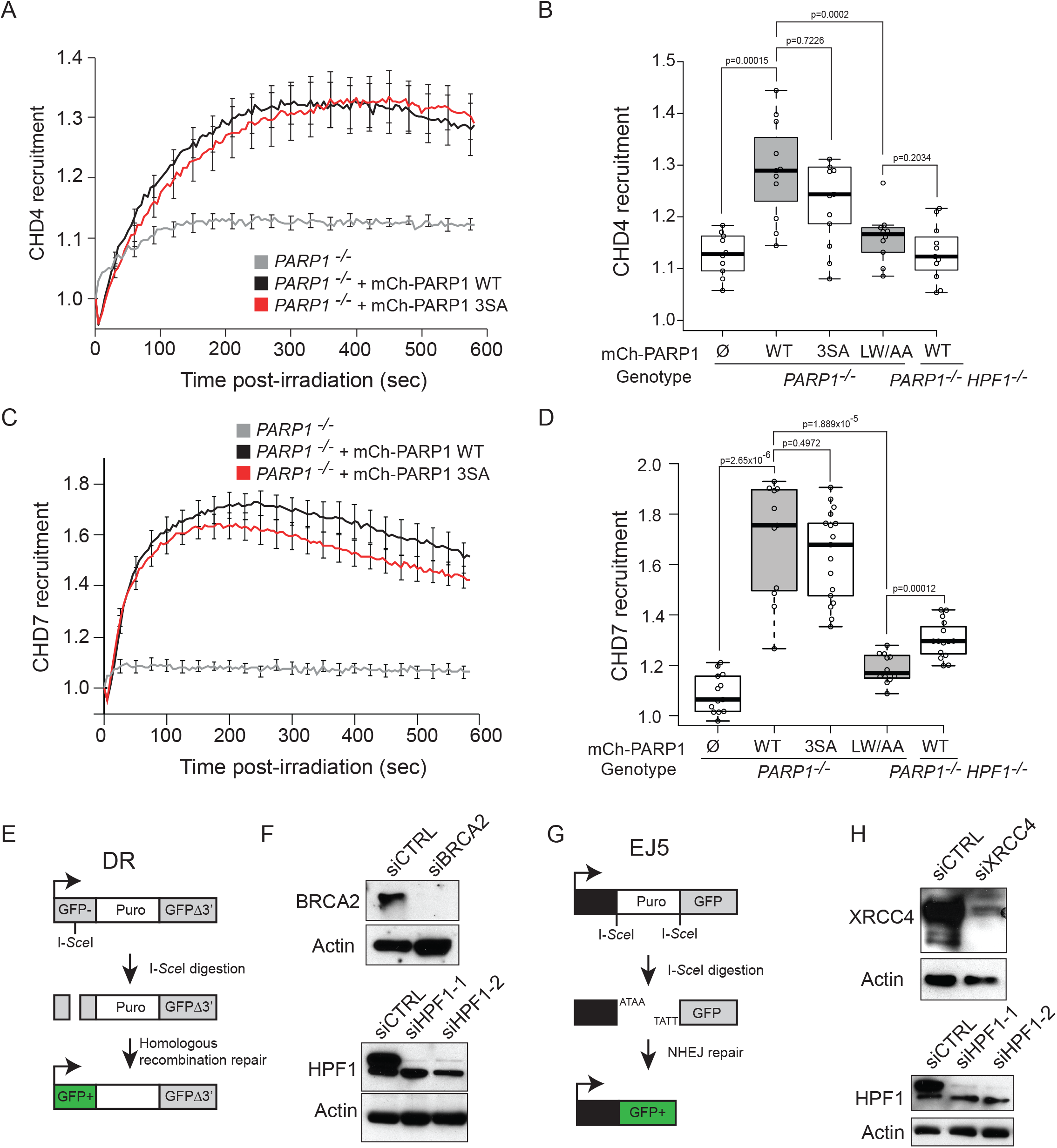
HPF1 contributes to efficient repair by homologous recombination and non-homologous end joining. (**A**) Recruitment kinetics of GFP-CHD4 to sites DNA damage in U2OS *WT*, *HPF1*^*KO*^ or *PARP1*^*KO*^ complemented or not with PARP1 WT or PARP1-3SA. (**B**) Quantification of mean intensity of GFP-CHD4 at sites DNA damage 200 s post-irradiation, in *PARP1KO* or *PARP1/HPF1* double knockout cells complemented or not with mCherry-PARP1 WT, PARP-3SA or PARP1-LW/AA mutants. ⵁ denotes no plasmid expression. (**C**) Recruitment kinetics of GFP-CHD7 to sites of DNA damage in U2OS *WT*, *HPF1*^*KO*^ or *PARP1*^*KO*^ complemented or not with PARP1 WT or PARP1-3SA. (**D**) Quantification of mean intensity of GFP-CHD7 at sites of DNA damage 200 s post-irradiation, in *PARP1KO* or *PARP1/HPF1* double knockout cells complemented or not with mCherry-PARP1 WT, PARP-3SA or PARP1-LW/AA mutants. ⵁ denotes no plasmid expression. (**E**) Schematic representation of the HR reporter assay (DR). After cleavage with I-SceI, the double-strand-breaks repaired by HR results in GFP expression. (**F**) Representative immunoblots showing the knockdown of BRCA2 and HPF1 in DR cells. Actin is used as a loading control. (**G**) Schematic representation of the NHEJ reporter assay (EJ5). Double cleavage by I-SceI removes the Puro cassette and the repair of the double-strand-break by NHEJ allows GFP expression. (**H**) Representative immunoblots showing knockdown of XRCC4 and HPF1 in EJ5 cells. Actin is used as a loading control.

## References

1. Kraus, W.L. & Hottiger, M.O. PARP-1 and gene regulation: progress and puzzles. Mol Aspects Med 34, 1109–23 (2013).

2. Ray Chaudhuri, A. & Nussenzweig, A. The multifaceted roles of PARP1 in DNA repair and chromatin remodelling. Nat Rev Mol Cell Biol (2017).

3. Eustermann, S. et al. Structural Basis of Detection and Signaling of DNA Single-Strand Breaks by Human PARP-1. Mol Cell 60, 742–754 (2015).

4. Ali, A.A.E. et al. The zinc-finger domains of PARP1 cooperate to recognize DNA strand breaks. Nat Struct Mol Biol 19, 685–692 (2012).

5. Langelier, M.-F., Planck, J.L., Roy, S. & Pascal, J.M. Structural basis for DNA damage-dependent poly(ADP-ribosyl)ation by human PARP-1. Science (New York, N.Y.) 336, 728–732 (2012).

6. Leidecker, O. et al. Serine is a new target residue for endogenous ADP-ribosylation on histones. Nat Chem Biol 12, 998–1000 (2016).

7. Buch-Larsen, S.C. et al. Mapping Physiological ADP-Ribosylation Using Activated Ion Electron Transfer Dissociation. Cell Reports 32, 108176 (2020).

8. Gibbs-Seymour, I., Fontana, P., Rack, J.G.M. & Ahel, I. HPF1/C4orf27 Is a PARP-1-Interacting Protein that Regulates PARP-1 ADP-Ribosylation Activity. Molecular cell 62, 432–442 (2016).

9. Suskiewicz, M.J. et al. HPF1 completes the PARP active site for DNA damage-induced ADP-ribosylation. Nature (2020).

10. Palazzo, L. et al. Serine is the major residue for ADP-ribosylation upon DNA damage. Elife 7 (2018).

11. Bonfiglio, J.J. et al. Serine ADP-Ribosylation Depends on HPF1. Molecular cell 65, 932–940.e6 (2017).

12. Rudolph, J., Roberts, G., Muthurajan, U.M. & Luger, K. HPF1 and nucleosomes mediate a dramatic switch in activity of PARP1 from polymerase to hydrolase. Elife 10 (2021).

13. Sun, F.H. et al. HPF1 remodels the active site of PARP1 to enable the serine ADP-ribosylation of histones. Nat Commun 12, 1028 (2021).

14. Mahadevan, J. et al. Q-FADD: A Mechanistic Approach for Modeling the Accumulation of Proteins at Sites of DNA Damage. Biophys J 116, 2224–2233 (2019).

15. Langelier, M.-F., Billur, R., Sverzhinsky, A., Black, B.E. & Pascal, J.M. HPF1 dynamically controls the PARP1/2 balance between initiating and elongating ADP-ribose modifications. bioRxiv, 2021.05.19.444852 (2021).

16. Prokhorova, E. et al. Serine-linked PARP1 auto-modification controls PARP inhibitor response. Nature Communications 12, 4055 (2021).

17. Juhász, S. et al. The chromatin remodeler ALC1 underlies resistance to PARP inhibitor treatment. Science Advances 6, eabb8626 (2020).

18. Shao, Z. et al. Clinical PARP inhibitors do not abrogate PARP1 exchange at DNA damage sites in vivo. Nucleic Acids Research 48, 9694–9709 (2020).

19. Gibson, B.A., Conrad, L.B., Huang, D. & Kraus, W.L. Generation and Characterization of Recombinant Antibody-like ADP-Ribose Binding Proteins. Biochemistry 56, 6305–6316 (2017).

20. Timinszky, G. et al. A macrodomain-containing histone rearranges chromatin upon sensing PARP1 activation. Nat Struct Mol Biol 16, 923–9 (2009).

21. Wang, Z. et al. Recognition of the iso-ADP-ribose moiety in poly(ADP-ribose) by WWE domains suggests a general mechanism for poly(ADP-ribosyl)ation-dependent ubiquitination. Genes Dev 26, 235–40 (2012).

22. Smith, R. et al. Poly(ADP-ribose)-dependent chromatin unfolding facilitates the association of DNA-binding proteins with DNA at sites of damage. Nucleic Acids Research 47, 11250–11267 (2019).

23. Sellou, H. et al. The poly(ADP-ribose)-dependent chromatin remodeler Alc1 induces local chromatin relaxation upon DNA damage. Molecular biology of the cell 27, 3791–3799 (2016).

24. Rother, M.B. et al. CHD7 and 53BP1 regulate distinct pathways for the re-ligation of DNA double-strand breaks. Nature Communications 11, 5775 (2020).

25. Smith, R., Sellou, H., Chapuis, C., Huet, S. & Timinszky, G. CHD3 and CHD4 recruitment and chromatin remodeling activity at DNA breaks is promoted by early poly(ADP-ribose)-dependent chromatin relaxation. Nucleic acids research 46, 6087–6098 (2018).

26. Pfeiffer, A. et al. Poly(ADP-ribosyl)ation temporally confines SUMO-dependent ataxin-3 recruitment to control DNA double-strand break repair. J Cell Sci 134 (2021).

27. Pan, M.R. et al. Chromodomain helicase DNA-binding protein 4 (CHD4) regulates homologous recombination DNA repair, and its deficiency sensitizes cells to poly(ADP-ribose) polymerase (PARP) inhibitor treatment. J Biol Chem 287, 6764–72 (2012).

28. Liu, C., Vyas, A., Kassab, M.A., Singh, A.K. & Yu, X. The role of poly ADP-ribosylation in the first wave of DNA damage response. Nucleic Acids Res 45, 8129–8141 (2017).

29. Barkauskaite, E., Jankevicius, G., Ladurner, A.G., Ahel, I. & Timinszky, G. The recognition and removal of cellular poly(ADP-ribose) signals. Febs j 280, 3491–507 (2013).

30. Murai, J. et al. Trapping of PARP1 and PARP2 by Clinical PARP Inhibitors. Cancer Res 72, 5588–99 (2012).

31. Poirier, G.G., de Murcia, G., Jongstra-Bilen, J., Niedergang, C. & Mandel, P. Poly(ADP-ribosyl)ation of polynucleosomes causes relaxation of chromatin structure. Proc Natl Acad Sci U S A 79, 3423–7 (1982).

32. Hananya, N., Daley, S.K., Bagert, J.D. & Muir, T.W. Synthesis of ADP-Ribosylated Histones Reveals Site-Specific Impacts on Chromatin Structure and Function. Journal of the American Chemical Society 143, 10847–10852 (2021).

33. de Murcia, G. et al. Modulation of chromatin superstructure induced by poly(ADP-ribose) synthesis and degradation. J Biol Chem 261, 7011–7 (1986).

34. Fontana, P. et al. Serine ADP-ribosylation reversal by the hydrolase ARH3. Elife 6 (2017).

35. Hatakeyama, K., Nemoto, Y., Ueda, K. & Hayaishi, O. Purification and characterization of poly(ADP-ribose) glycohydrolase. Different modes of action on large and small poly(ADP-ribose). J Biol Chem 261, 14902–11 (1986).

36. Luijsterburg, M.S. et al. PARP1 Links CHD2-Mediated Chromatin Expansion and H3.3 Deposition to DNA Repair by Non-homologous End-Joining. Mol Cell 61, 547–562 (2016).

37. Bacic, L. et al. Structure and dynamics of the chromatin remodeler ALC1 bound to a PARylated nucleosome. bioRxiv, 2021.06.18.448936 (2021).

38. Beaudouin, J., Mora-Bermúdez, F., Klee, T., Daigle, N. & Ellenberg, J. Dissecting the Contribution of Diffusion and Interactions to the Mobility of Nuclear Proteins. Biophys J 90, 1878–94 (2006).

39. Polo, S.E., Kaidi, A., Baskcomb, L., Galanty, Y. & Jackson, S.P. Regulation of DNA-damage responses and cell-cycle progression by the chromatin remodelling factor CHD4. Embo j 29, 3130–9 (2010).

40. Richardson, C., Moynahan, M.E. & Jasin, M. Double-strand break repair by interchromosomal recombination: suppression of chromosomal translocations. Genes Dev 12, 3831–42 (1998).

41. Czarna, A. et al. Structures of Drosophila cryptochrome and mouse cryptochrome1 provide insight into circadian function. Cell 153, 1394–405 (2013).

42. Gunn, A. & Stark, J.M. I-SceI-based assays to examine distinct repair outcomes of mammalian chromosomal double strand breaks. Methods Mol Biol 920, 379–91 (2012).

